# RNPS1 in PSAP complex controls periodic pre-mRNA splicing over the cell cycle

**DOI:** 10.1101/2023.11.20.567984

**Authors:** Kazuhiro Fukumura, Akio Masuda, Jun-ichi Takeda, Osamu Nagano, Hideyuki Saya, Kinji Ohno, Akila Mayeda

## Abstract

Cell cycle progression requires periodic gene expression through splicing control. However, the splicing factor that directly controls this cell cycle-dependent splicing remain unknown. Cell cycle-dependent expression of the *AURKB* (aurora kinase B) gene is essential for chromosome segregation and cytokinesis. We previously reported that RNPS1 is essential to maintain precise splicing in *AURKB* intron 5. Here we show that RNPS1 plays this role in PSAP complex with PNN and SAP18, but not ASAP complex with ACIN1 and SAP18. Whole-transcriptome sequencing of RNPS1- and PNN-deficient cells indicated that RNPS1, either alone or as PSAP complex, is an essential splicing factor for a subset of introns. Remarkably, protein expression of RNPS1, but not PNN, is coordinated with cyclical splicing in PSAP-controlled introns including *AURKB* intron 5. The ubiquitin-proteasome pathway is involved in the periodic decrease of RNPS1 protein level. RNPS1 is a key factor that controls periodic splicing during the cell cycle.

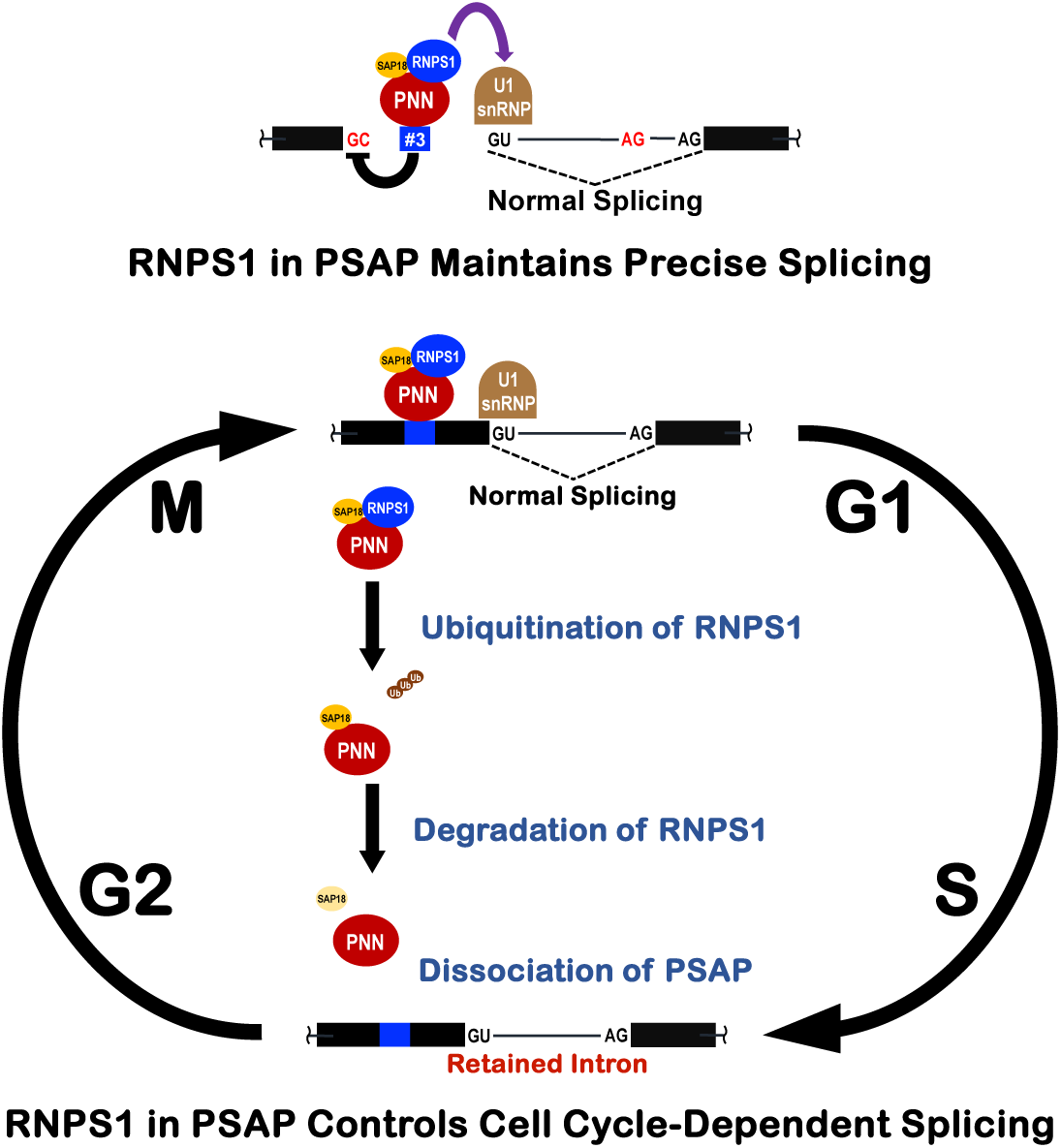

**Highlights:**

- The *AURKB* gene encodes a protein kinase essential for cytokinesis and cell cycle
- PSAP complex (RNPS1, PNN, SAP18) is required for precise splicing in *AURKB* intron 5
- RNPS1 protein in PSAP complex controls periodical splicing during the cell cycle
- Cyclical decrease of RNPS1 protein is mediated by the ubiquitin-proteasome pathway

## INTRODUCTION

Human pre-mRNA splicing takes place in the huge multimolecular complex, termed spliceosome, that is a highly organized dynamic ribonucleoprotein machine comprised of five snRNPs and numerous proteins (reviewed in Ref.^1,2^). Amidst the panoply of proteins in this spliceosome, it is amazing to observe that each individual protein factor often drastically changes the outcome of splicing in a precisely regulated manner.

We originally identified human RNPS1 (RNA binding protein with serine-rich domain 1) as a general splicing activator *in vitro*^3^. We next found that RNPS1 exists in the spliceosome and regulates a variety of alternative splicing events, often collaborating with other splicing regulators including SRSF11 (p54), TRA2B (hTra2ϕ3), and PNN (PININ)^4,5^. RNPS1 was also identified as a peripheral factor of the exon junction complex (EJC) formed on spliced mRNA (reviewed in Ref.^6,7^).

ACIN1 (ACINUS) and PNN are scaffold proteins and form two alternative ternary complexes with RNPS1 and SAP18, termed ASAP and PSAP, respectively^8,9^. ACIN1 is involved in apoptosis and regulation of splicing and transcription^10–12^ and PNN was originally identified as a desmosome-associated protein but also function as a regulator of alternative splicing^13,14^.

Since ACIN1 bridges the ASAP complex to the EJC core^14^ and PNN in PSAP complex is structurally related to ACIN1^9^, both ASAP and PSAP complexes must interact with the EJC. Recent transcriptome-wide analysis indicated that the ASAP and PSAP complexes regulate distinct alternative splicing either in EJC core-dependent or -independent manner^14^. On the other hand, the EJC core alone, without any peripheral factors repress aberrant re-splicing on mature mRNA in cancer cells, implicating an important role in terminating splicing^15^. Many lines of evidence support that ASAP and PSAP complexes, with or without EJC core, are general regulators of splicing (reviewed in Ref.^7^). However, their individual biological and physiological significance remains to be investigated.

We have been studying splicing regulation involved in mitotic cell-cycle progression using the *AURKB* (aurora kinase B) gene as a model. The *AURKB* gene encodes a serine/threonine protein kinase that is essential in chromosome segregation and cytokinesis during the cell cycle (reviewed in Ref.^16^). We previously showed that RNPS1, recruited by the EJC core, is critical for efficient and faithful splicing in mitotic cell cycle-related genes including *AURKB*^17^. Indeed, RNPS1 depletion causes a reduction of the AURKB protein due to multiple aberrant splicing, resulting in abnormal multinucleation phenotype that is fully rescued by ectopic expression of *AURKB*^18^. Regarding the mechanism of action, we found that RNPS1 protein binds directly to a specific element in exon 5 upstream of the authentic 5′ splice site, and this binding is critical to prevent multiple aberrant splicing using a common pseudo 5′ splice site^18^. Here we demonstrate that RNPS1 plays a role as the PSAP component in preventing aberrant splicing of *AURKB* pre-mRNA.

It was reported that SR protein kinase CLK1 (CDC-like kinase 1) controls many periodically spliced pre-mRNAs, including *AURKB*, during the cell cycle^19^. However, splicing factors were not known to be directly involved in cyclical splicing. We have discovered that the protein level of RNPS1 is coordinated with cell cycle-dependent periodic splicing of *AURKB* pre-mRNA. Here we propose a novel function of RNPS1 in PSAP complex, as a critical modulator that temporally controls periodic splicing observed during cell cycle.

## RESULTS

### PSAP promotes efficient and precise splicing of *AURKB* pre-mRNA

Previously, we reported that siRNA-mediated knockdown of RNPS1 induced aberrant splicing in the *AURKB* intron 5 and *MDM2* intron 10 (Figure S1A), and we identified the functional RNPS1 binding site upstream of the *AURKB* intron 5^18^. On the other hand, our previous yeast two-hybrid screening revealed that RNPS1 can bind to ACIN1, PNN, GPATCH8 and LUC7L3 (hLuc7A)^4^. This observation is consistent with the finding that ACIN1 and PNN form the ASAP and PSAP complex, respectively, with the common components RNPS1 and SAP18^8,9^.

We then performed knockdown of all the components of the PSAP and ASAP complexes (RNPS1, ACIN1, PNN and SAP18) in HeLa cells. We confirmed that each siRNA led to effective depletion of the target mRNA and the protein product (Figures 1A, B). Notably, the knockdown of RNPS1, ACIN1 and PNN also caused co-depletion of SAP18 protein (Figure 1A), however SAP18 mRNA levels were barely changed (Figure 1B), indicating that SAP18 protein is stabilized in either ASAP or PSAP complex. The co-depletion of SAP18 protein was also observed by the knockdown of RNPS1, ACIN1 and PNN in HEK293 cells (Figure S2A; Cf. Figure S2B)

**Figure 1.**
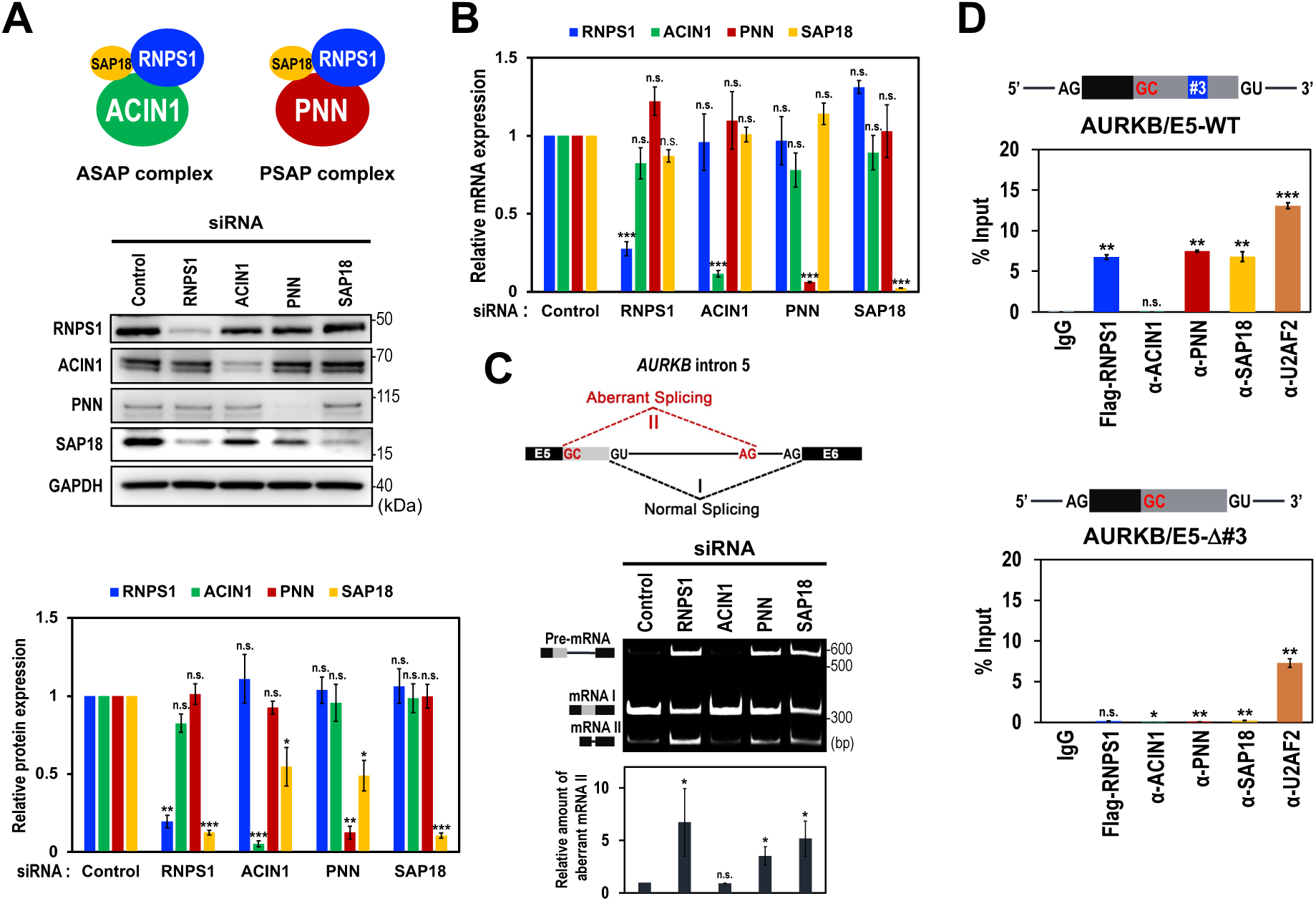
PSAP component binds to a specific site that promotes normal splicing of the *AURKB* pre-mRNA. (A) Schematic structures of ASAP and PSAP complexes (upper scheme). siRNA-mediated depletion of the indicated ASAP/PSAP components in HeLa cells were analyzed by Western blotting with indicated antibodies (middle panel). Individual bands on the Western blots were quantified and the relative values were standardized to that in the control siRNA (lower graph). Means ± standard errors (SE) are given for three independent experiments and Welch’s t-test values were calculated (*p < 0.05, **p < 0.005, ***p < 0.0005, n.s. p > 0.05). See also Figure S2A. (B) The mRNA levels of ASAP/PSAP components in (A) were analyzed by RT-qPCR using specific primer sets. See (A) for the statistical analysis. See also Figure S2B. (C) Splicing defect and aberrant splicing in the endogenous *AURKB* gene, induced by siRNA-mediated depletion of PSAP proteins, were detected by RT–PCR, visualized by PAGE, and individual bands on the PAGE gel were quantified. The ratios of the aberrantly spliced mRNA value (II) to the sum of the spliced mRNAs value (I+II) were standardized to that of the control siRNA and plotted. See (A) for the statistical analysis. See also Figures S1, S3. (D) RNA immunoprecipitation assay was performed using HeLa cells co-transfected with Flag-RNPS1 expression plasmids and the indicated reporter AURKB-Exon 5 mini-genes. After Immunoprecipitation using the indicated antibodies, immunoprecipitated RNAs were quantified by RT–qPCR using specific primers. See (A) for the statistical analysis.

Then, we examined whether these PSAP- and ASAP-components operate together with RNPS1 to ensure precise splicing in the *AURKB* intron 5 (Figure 1C) and the *MDM2* intron 10 (Figure S1B). We found that the knockdown of PNN and SAP18, as well as RNPS1, generated unspliced pre-mRNA and induced aberrant splicing at expense of normal mRNA, but knockdown of ACIN1 did not. This confirms that the PSAP, but not ASAP, plays a role to restore the normal splicing in *AURKB* intron 5. The knockdown of other RNPS1-binding proteins, GPATCH8 and LUC7L3, barely affected in normal splicing (I) of the *AURKB* intron 5 and *MDM2* intron 10 (Figure S1B).

Previously, we demonstrated that RNPS1 binds to a specific *cis-*acting element (termed #3) located upstream of the authentic 5′ splice site of intron 5 (Figure 1D, upper scheme)^18^. We thus examined whether the PSAP complex binds to this *cis*-acting element by RNA immunoprecipitation experiments using antibodies against each of the PSAP components (Figure 1D, lower graphs). As we expected, RNPS1, PNN and SAP18 bound to the AURKB/E5-WT, but ACIN1 did not. Notably, deletion of this *cis*-acting element ‘#3’ greatly reduced the binding of the PSAP components. We conclude that PSAP binding to the specific element ‘#3’ is critical for efficient and precise splicing in the *AURKB* intron 5.

### PSAP either represses the pseudo 5′ splice site or activates the authentic 5′ splice site in *AURKB* pre-mRNA

Since the ‘#3’ element is located between the upstream pseudo 5′ splice site and the downstream authentic 5′ splice site, we postulated two conceivable mechanisms leading to the normal splicing; either (i) PSAP represses the pseudo 5′ splice site, and/or (ii) PSAP activates the authentic 5′ splice site (Figure S3A). To examine distance-dependent effects of PSAP binding, such as interaction with and/or steric hindrance to the U1 snRNP, we increased the distance of the ‘#3’ element from either the authentic 5′ splice site or the pseudo 5′ splice site (Figure S3B).

These two variant *AURKB* pre-mRNAs, as well as the original *AURKB* pre-mRNAs, had equivalent repression of the normal 5′ splice site by either RNPS1-knockdown or PNN-knockdown (Figure S3C), suggesting that both postulated mechanisms (see above) are involved. Several previous studies indeed support both possibilities (see DISCUSSION).

### Individual subsets of introns are spliced out by RNPS1, PNN and PSAP

To test whether PSAP might have a general role in controlling splicing, we performed siRNA-mediated knockdown of RNPS1 and PSAP-specific component PNN in HEK293 cells followed by the whole-transcriptome sequencing (RNA-Seq). The sequencing reads were mapped to the human genome sequences and we analyzed splicing patterns, both negatively and positively changed, in five major classes (Figure S4, Tables S2, S3).

The most frequent class of the overlapping events was intron retention type (131/289; Figure S4, Table S4), and remarkably, these are all intron retention promoted, not repressed (Figure 2A), indicating that these 131 introns are PSAP-dependently spliced. The data also reveal that RNPS1 and PNN solely (not in PSAP complex) promote splicing in 791 and 53 introns, respectively (Figure 2A).

**Figure 2.**
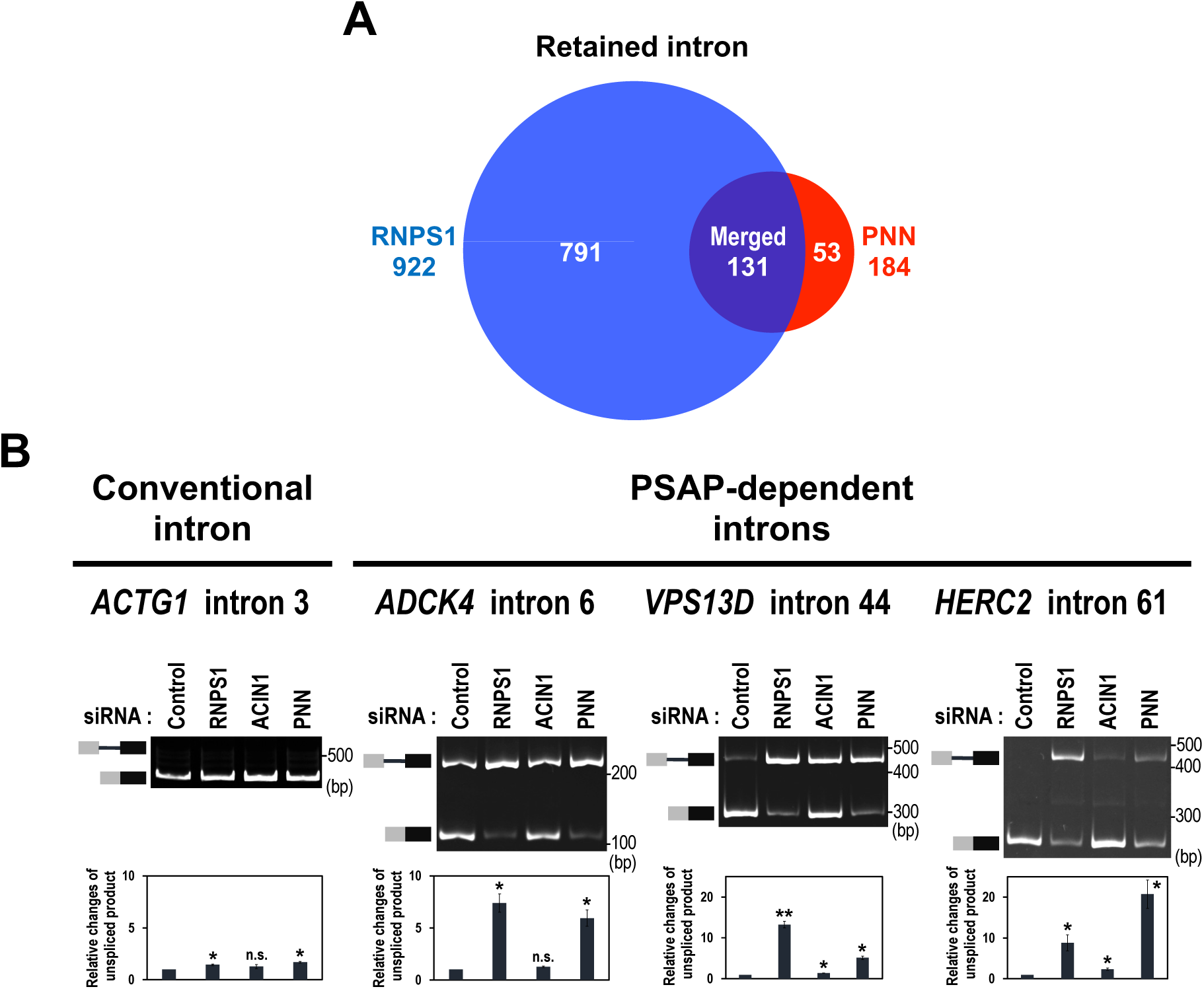
RNPS1 and PNN are general splicing factors for individual subsets of introns. (A) Venn diagram of retained introns that are generated by RNPS1-and PNN-knockdown HEK293 cells. See also Figure S4 and Tables S2–S4. (B) *In cellulo* splicing assays of the pre-mRNAs including a conventional intron and three representative PSAP-dependent introns. After the indicated siRNA-mediated knockdown in HEK293 cells, indicated endogenous splicing was analyzed by RT–PCR followed by PAGE (upper panel). The unspliced RNA products were quantitated by RT–qPCR and the relative values were standardized to that in the control siRNA (lower graph). Means ± SE are given for three independent experiments and Welch’s t-test values were calculated (*p < 0.05, **p < 0.005, n.s. p > 0.05).

We validated the PSAP-dependent splicing defect by RT–PCR with three representative PSAP-dependent introns (arbitrarily chosen from 131 introns in Table S2). By depletion of PSAP components (RNPS1 and PNN), we observed unspliced pre-mRNAs with all the PSAP-dependent introns, whereas the control authentic *ACTG1* intron 3 was fully spliced out (Figure 2B). These results implicate that PSAP promotes splicing in a subset of introns.

### PSAP-controlled introns are periodically spliced over the cell cycle

A previous RNA-Seq analysis at different stages of the cell cycle identified ∼1300 genes with changes of cell cycle-dependent alternative splicing, and notably, the intron retention event of *AURKB* intron 5 was characterized as a representative model^19^. It was reported that the excision of *AURKB* intron 5 is periodically modulated and this splicing activity reaches the maximum around M phase (Figure 3A). Using the same *AURKB* pre-mRNA including intron 5, we demonstrated that RNPS1 in PSAP complex binds upstream of the 5′ splice site and promotes its precise splicing (Figures 1C, D)^18^. We thus hypothesized that PSAP also plays a role in cell cycle-dependent splicing observed in *AURKB* intron 5.

**Figure 3.**
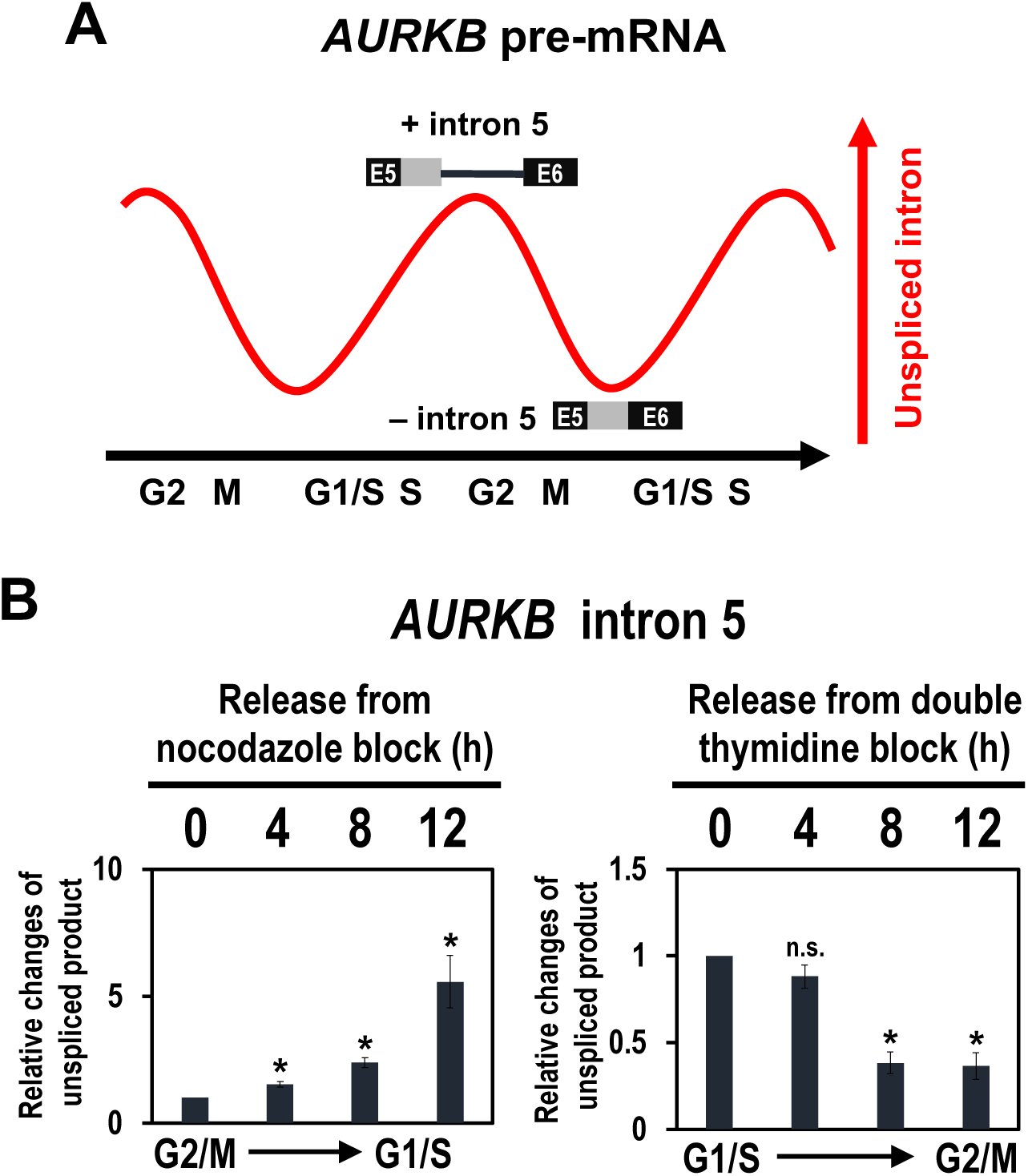
Reported periodic *AURKB* pre-mRNA splicing is recapitulated in synchronized cells during the cell cycle. (A) Schematic representation of cyclical splicing in *AURKB* intron 5 (modified from Ref.^19^). (B) HEK293 cells were synchronized in G2/M phase by treating with nocodazole and synchronized in G1/S phase by treating with double thymidine. Cells were washed, released, and harvested at the indicating time points. Endogenous splicing in the *AURKB* intron 5 was analyzed by RT–qPCR and the unspliced products were standardized to that at the start time, 0 h. Means ± SE are given for three independent experiments and Welch’s t-test values were calculated (*p < 0.05, n.s. p > 0.05). See also Figure S5, S6.

We first recapitulated the periodic splicing of *AURKB* intron 5 during cell cycle progression with HEK293 cells (Figure 3B). We treated the cells with the microtubule-destabilizing reagent, nocodazole, which arrests cells cycle at the start of the G2/M phase (nocodazole release 0 h). Using RT–PCR followed by PAGE analysis and quantitative RT–PCR (RT– qPCR), we observed increasing unspliced intron 5 from G2/M to G1/S phase (Figure S5, Figure 3B). Whereas from G1/S to G2/M phase, we found decreasing unspliced intron 5 in synchronized HEK293 cells after release from the double thymidine block at the G1/S boundary (Figure 3C).

It is important to confirm whether the observed cyclical splicing modulation is a general event in the PSAP-controlled introns. Using three representative PSAP-dependently spliced introns (Figure 2B), we examined the splicing changes during the cell cycle using nocodazole and double thymidine to synchronize HEK293 cells (Figure 4, Figure S5). Remarkably in these PSAP-controlled introns, we also observed periodic changes of unspliced introns as observed in *AURKB* intron 5 pre-mRNA (Figure 4, Cf. Figure 3B). These periodic patterns are evidently unique when compared with those observed in a conventional intron. Our results thus provided the mechanistic basis underlying the reported periodical pattern of splicing activity (Figure 3A). Here we propose that PSAP promotes periodic splicing linked to the cell cycle.

**Figure 4.**
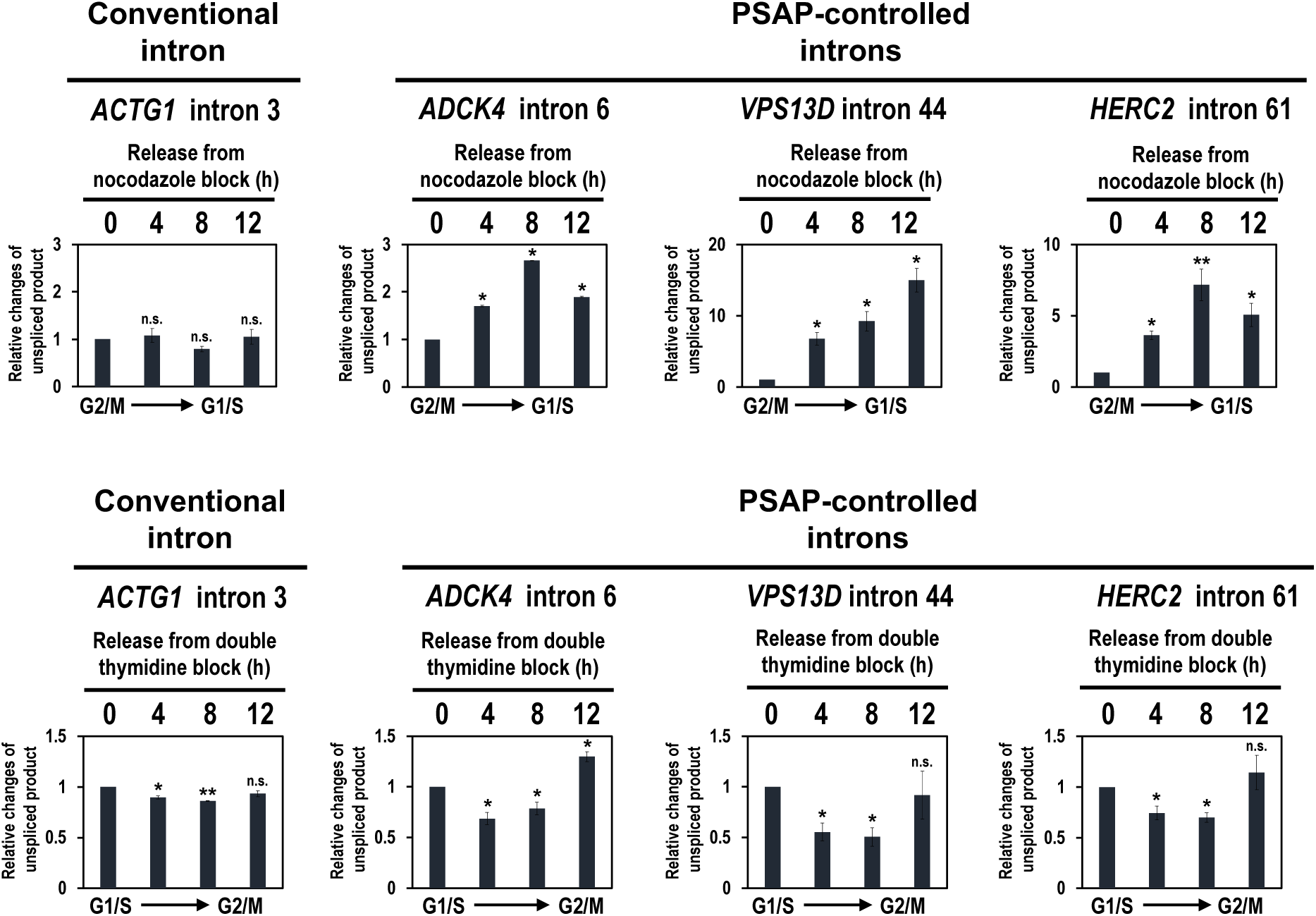
Periodic modulation of unspliced introns across the cell cycle was observed in PSAP-controlled introns but not in conventional intron. HEK293 cells were synchronized in G2/M phase by nocodazole block (upper panels) and synchronized in G1/S phase by double thymidine block (lower panels). Endogenous splicing in the indicated conventional intron and PSAP-controlled introns was analyzed by RT–qPCR. Quantification and the statistical analysis were performed as described in Figure 3B (*p < 0.05, **p < 0.005, n.s. p > 0.05). See also Figure S5, S6.

### RNPS1 in PSAP controls periodic splicing over the cell cycle

We assumed that PSAP-controlled periodic splicing was modulated by the expression level changes in any of the PSAP component. We thus examined the protein levels of PSAP components during the cell cycle using nocodazole synchronized HEK293 cell (Figure 5A). Intriguingly, RNPS1 protein level was at a maximum in G2/M phase (nocodazole release 0 h) and decreased through cell cycle progression (nocodazole release 4–8 h). The decrease in RNPS1 protein was coordinated with the level of AURKB protein, which was at a high level in G2/M and declined to a low level in G1/S (Figure 5A)^20^. In contrast, protein levels of other PSAP components PNN and SAP18, as well as an ASAP component ACIN1, were not significantly changed. Notably, we did not observe such a change in RNPS1 mRNA expression level during cell cycle (Figure 5B) as previously reported in human RNPS1 (E5.1)^21^. These data together indicate that the reduction of RNPS1 protein level is regulated at the protein level, but not at the mRNA level.

**Figure 5.**
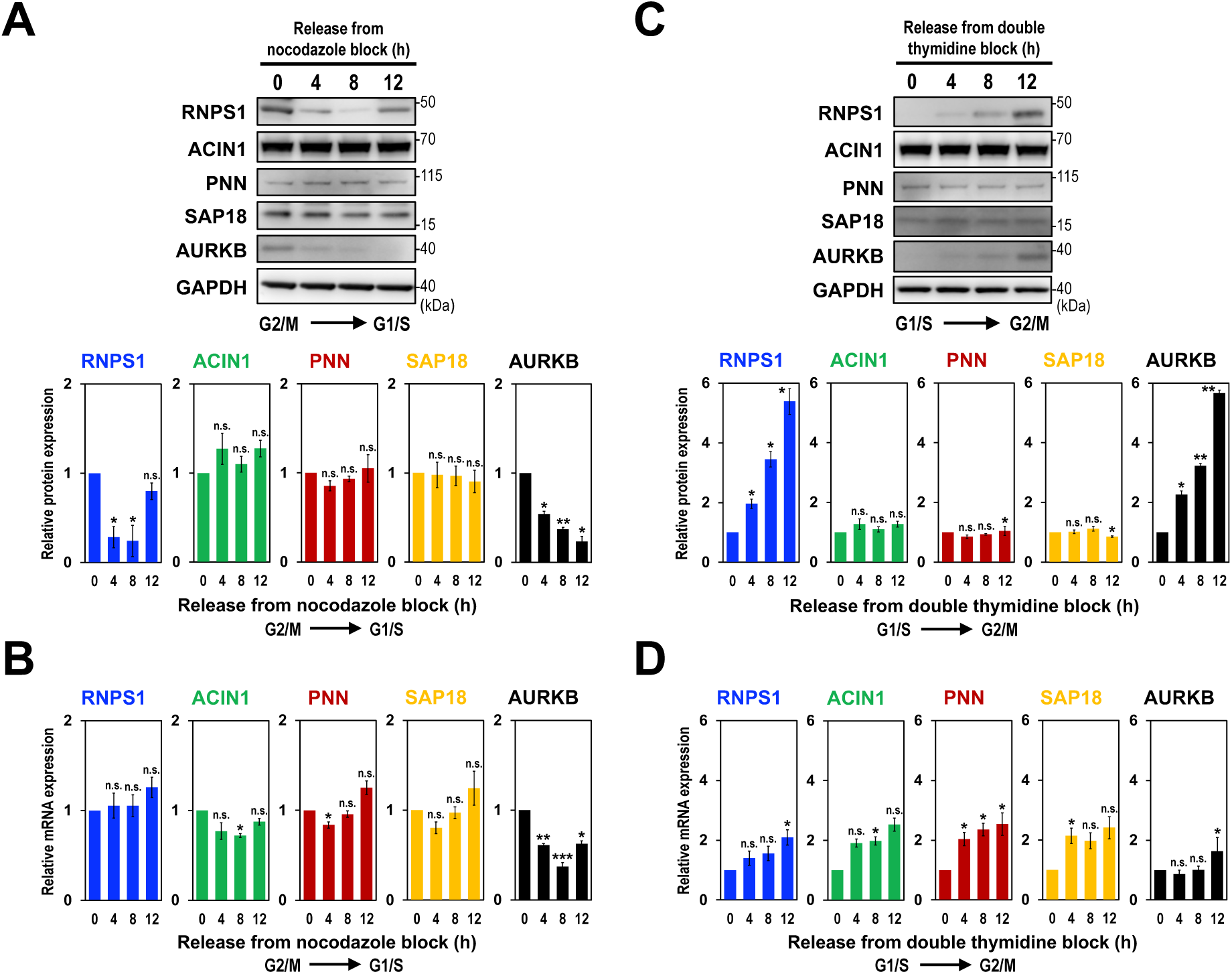
The level of RNPS1 protein, but not other ASAP/PSAP proteins, is coordinated with AURKB protein level during the cell cycle. (A) HEK293 cells were synchronized in G2/M phase by nocodazole block and released cells were harvested at indicated time points. ASAP/PSAP proteins were analyzed by Western blotting with indicated antibodies (upper panel). Individual bands on the Western blots were quantified and the relative values were standardized to that at the start time, 0 h (lower graph). Means ± SE are given for three independent experiments and Welch’s t-test values were calculated (*p < 0.05, **p < 0.005, ***p < 0.0005, n.s. p > 0.05). See also Figure S6. (B) mRNA levels in (A) were analyzed by RT–qPCR and the relative values were standardized to that at the start time, 0 h. See (A) for the statistical analysis. (C) HEK293 cells were synchronized at G1/S phase by double thymidine block and ASAP/PSAP proteins were analyzed as described in (A). See (A) for the statistical analysis. See also Figure S6. (D) mRNA levels in (C) were analyzed by RT–qPCR and the relative values were standardized to that at the start time, 0 h. See (A) for the statistical analysis.

We also examined the levels of PSAP components for the latter part of the cell cycle using the double thymidine block synchronized HEK293 cells (Figure 5C). We found that RNPS1 protein was minimal at G1/S phase (double thymidine release 0 h) and greatly increased as the cell cycle progressed to G2/M phase (double thymidine block 4–12 h), which was well coordinated again with the level of AURKB protein. The RNPS1 protein increase could be due to the mRNA level in which RNPS1 was increased, together with AURKB, from G1/S into G2/M phase (Figure 5D).

Since RNPS1 and AURKB protein expressions were synchronized during the cell cycle (Figures 5A, C), we assumed that the splicing activator RNPS1 was responsible for the observed cell cycle-dependent splicing modulation (Figures 3, 4). To test this assumption, we performed nocodazole treatment in RNPS1-knockdown HEK293 cells, and analyzed splicing changes in the two PSAP-controlled introns (Figure S6A). Even at the RNPS1-upregulated G2/M phase, we observed that RNPS1-knockdown generated the unspliced product. Nocodazole treatment in PNN-knockdown HEK 293 cells also increased the unspliced product (Figure S6B) confirmed that RNPS1, as a component of the PSAP, promotes the periodic pre-mRNA splicing linked to cell cycle progression.

### Periodic decrease of RNPS1 protein is mediated by the ubiquitin-proteasome pathway

Ubiquitin-mediated proteolysis is one of the keys to regulating cell cycle progression^22^. RNPS1 is indeed polyubiquitinated by K48-linkage and degraded by the proteasome^23^. We assumed that periodical degradation of RNPS1 protein is controlled by ubiquitin-proteasome pathway.

We treated HEK293 cells with proteasome inhibitor MG132 and we examined the protein level of PSAP components. As we expected, RNPS1 protein was stabilized in the presence of MG132 whereas the level of other ASAP/PSAP proteins, ACIN1, PNN and SAP18, were not changed at all (Figure 6A). Since the degradation of RNPS1 causes co-destabilization of SAP18 (Figure 1A), the functional PSAP complex appears to be dissociated, that is the cause of generated unspliced intron . Taken together, we conclude that ubiquitin-proteasome pathway plays a key role in the periodic fluctuation of RNPS1 protein level, leading to observed periodic splicing in accordance with the cell cycle (Figure 6B).

**Figure 6.**
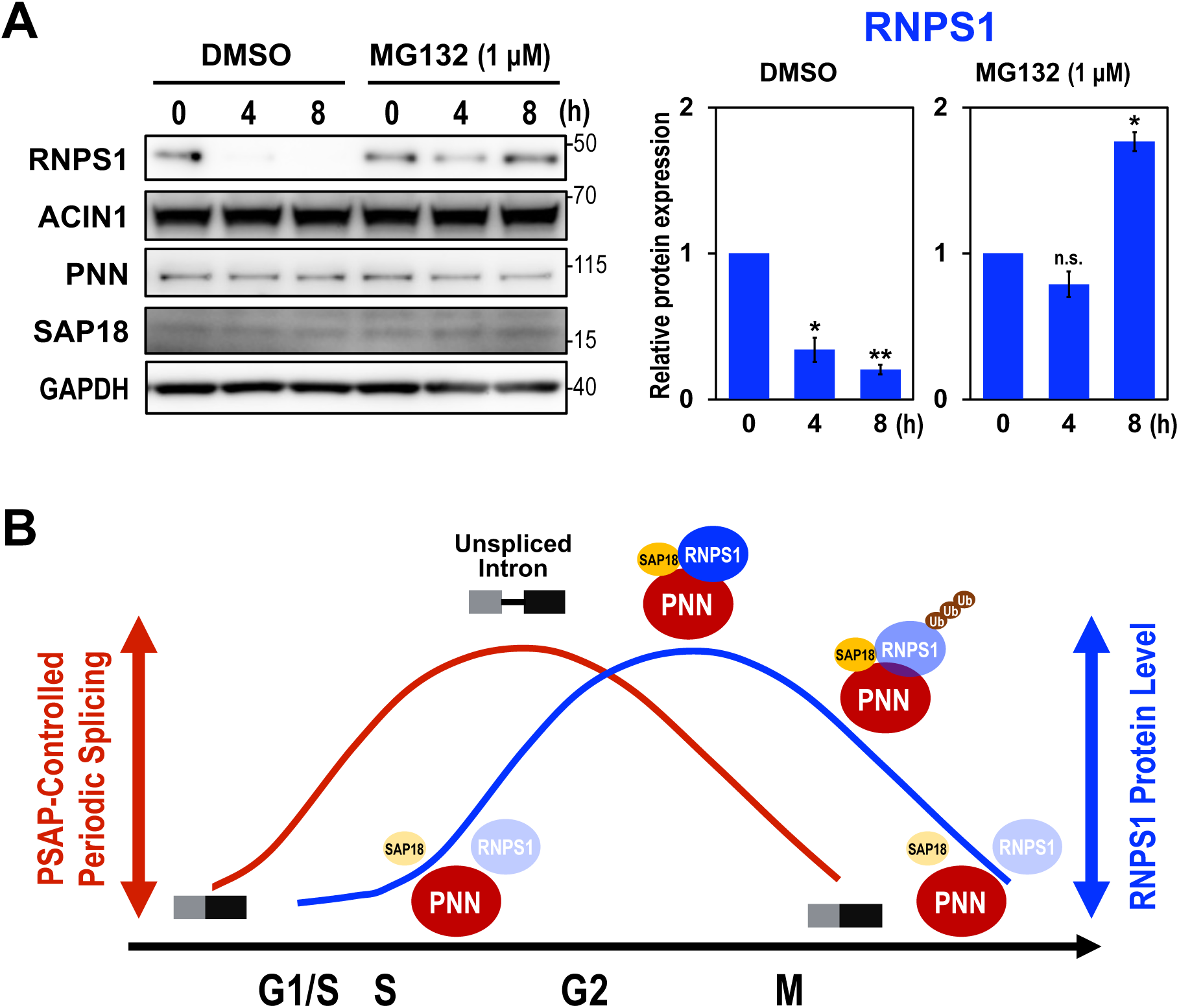
PSAP-mediated cyclical splicing is controlled by RNPS1 protein level through the ubiquitin-proteasome pathway. (A) HEK293 cells were cultured for 4 h and 8 h with proteasome inhibitor MG132 and the protein levels were analyzed by Western blotting using the antibody against each ASAP/PSAP protein. A common solvent, dimethyl sulfoxide (DMSO), was added to the medium (final concentration at 0.1%). The RNPS1 band on the Western blots were quantified and the relative values were standardized to that at the start time, 0 h (right graph). Means ± SE are given for three independent experiments and Welch’s t-test values were calculated (*p < 0.05, **p < 0.005, n.s. p > 0.05). (B) A schematic model of RNPS1-mediated periodic splicing during cell cycle. RNPS1 protein, but neither PNN nor SAP18 protein, in PSAP plays an essential role to control periodic splicing over the cell cycle progression. The ubiquitin-mediated proteolysis of RNPS1 also destabilizes SAP18 that induces dissociation of PSAP complex, leading to the splicing defect.

## DISCUSSION

### The PSAP complex is critical to maintain splicing fidelity

We previously reported that RNPS1 interacts with a specific *cis*-element and ensures precise splicing in the *AURKB* pre-mRNA, which is essential for chromosome segregation and cytokinesis^18^. Here, we demonstrate that RNPS1, as a component of PSAP complex with PNN and SAP18, binds to the same *cis*-element in the upstream exon, leading to the precise splicing on *AURKB* intron 5 (Graphical Abstract, upper panel).

The PSAP components (PNN, SAP18 and RNPS1) are all associated with the EJC core as peripheral factors (reviewed in Ref.^6,7^). Therefore, it is conceivable that the EJC recruits the PSAP complex on the target pre-mRNA and PSAP itself functions to promote precise splicing. Indeed, we previously found that EJC core on pre-mRNA recruits RNPS1 as a key factor to promote splicing involved in mitotic cell cycle^17^.

Regarding the mechanism of PSAP-mediated control to ensure precise splicing in *AURKB* pre-mRNA, our data implicate that it occurs either through repression of the upstream pseudo 5′ splice site (without direct masking) or through activation of the downstream authentic 5′ splice site, likely through the interaction with U1 snRNP (Figure S3A; see ‘Limitations of the study’). Since the downstream authentic 5′ splice site is much weaker than the upstream pseudo 5′ splice site^18^, the intrinsic RNPS1 activity as a general splicing activator^3^, in a component of PSAP, must be a prerequisite for the use of the weak authentic 5′ splice site. On the other hand, repression of the pseudo 5′ splice site cannot be ruled out because RNPS1-knockdown produces five aberrant mRNAs using one common upstream pseudo 5′ splice site and the downstream five 3′ splice sites in endogenous *AURKB* pre-mRNA^18^. This ‘repression’ scenario is consistent with two recent observations that EJC-recruited PSAP represses distant pseudo and recursive 5′ splice site^24,25^, however, the underlying mechanism to repress such distant 5′ splice site without a steric hindrance remains unknown.

### RNPS1 functions in periodical splicing during the cell cycle

Cyclical removal of *AURKB* intron 5 contributes to periodic AURKB protein expression during the mitotic cell cycle^19^; however, the involved mechanism remains to be elucidated. Our finding could provide a breakthrough in addressing this important open question. Here we propose that splicing activator RNPS1 in PSAP complex not only functions to ensure precise splicing, but also plays an essential role in controlling periodic splicing linked to cell cycle (Graphical Abstract, lower panel).

Our RNA-Seq analysis in RNPS1- and PNN-knockdown cells showed that most affected splicing events were intron retentions, and we could validate that these intron retention events were all induced from G2/M phase to G1/S phase as observed previously in *AURKB* intron 5^19^. We eventually discovered that the level of RNPS1 protein is down-regulated in G1/S phase by the ubiquitin proteasome pathway and it is gradually up-regulated to its maximum level at M phase. This characteristic temporal fluctuation of the RNPS1 protein level is well accounted for by the periodic splicing in accordance with the cell cycle (Figure 6B). Since the protein levels of other PSAP components, PNN and SAP18, remain constant during the cell cycle, it is likely that the functional PSAP complex is dissociated by the lack of RNPS1 component.

Previously, it was reported that knockdown of EJC-core component, RBM8A (Y14), induces abnormal nuclear structure, multinucleated cells, G2/M cell cycle arrest, and genome instability (Ref.^17,26^; and references therein), which is a similar phenotype to that caused by the knockdown of PSAP component, RNPS1^18^. In mouse models, it was reported that haploinsufficiency of MAGOH, the other EJC core component, causes the defect of mitosis of neural stem cells, ending up in microcephaly^27^. Together, it is very likely that the EJC recruits the functional PSAP complex on a subset of pre-mRNAs to control periodic splicing during mitotic cell cycle, as well as to ensure efficient and precise splicing at least in *AURKB* pre-mRNA.

We demonstrate that PSAP, but not ASAP, controls the faithful splicing and alternative splicing of transcripts involved in the mitotic cell cycle. Therefore, the function of these two complexes is determined by only one distinct component of PSAP and ASAP, i.e., PNN and ACIN1 (Figure 1A, top schemes). Consistently, a recent report showed that PSAP and ASAP form functionally distinct complexes with the EJC core to confer distinct alternative splicing activities^14^. On the other hand, since RNPS1 is a common component of PSAP and ASAP, periodic changes of RNPS1 protein would also be expected to modulate the function of ASAP during the cell cycle. Indeed, a possible function of ASAP in the cell cycle is suggested by the fact that depletion of the component of ASAP, ACIN1, cause defects in cell cycle progression^28^. Moreover, the genome-wide identification of RNA targets suggests the role of ACIN1 in cell cycle^12^. Taken together, we speculate that PSAP and ASAP complexes may target distinct subsets of pre-mRNAs and control periodic changes of splicing during the cell cycle. Although it is out of scope for this article, there is considerable interest in identifying these distinct subsets of pre-mRNAs.

### Limitations of the study

We previously found that RNPS1 can bind to a component of U1 snRNP, LUC7L3 (one of the human paralogs of yeast Luc7A)^4^. It was also shown that LUC7L3 overexpression changes the selection of the alternative 5′ splice site *in cellulo*^29^. Moreover, recent study showed that three human paralogs, LUC7L, LUC7L2 and LUC7L3, as components of the U1 snRNP, regulates unique alternative splicing profiles^30^.

We thus tested our hypothesis that the weak authentic 5′ splice site is activated through RNPS1 binding to LUC7L3 in the U1 snRNP. However, depletion of LUC7L3 has no effects on the PSAP-dependent *AURKB* and *MDM2* pre-mRNA splicing (Figure S1B), suggesting that RNPS1–LUC7L3 interaction is not involved in these two PSAP-dependent splicing events. Of course, we cannot rule out the possibility of RNPS1 interaction with other U1 snRNP components.

Further studies are needed to elucidate the PSAP-mediate molecular mechanism to ensure splicing fidelity observed in *AURKB* and *MDM2* pre-mRNAs.

## MATERIALS AND METHODS

### Human cell line

HeLa and HEK293 cells (ATCC) were cultured in Dulbecco’s modified Eagle’s medium (DMEM; Wako) supplemented with 10% fetal bovine serum (Sigma-Aldrich) and 1% penicillin-streptomycin (Nakarai Tesque). Cells were grown at 37°C in 5% CO_2_.

### Construction of expression plasmids

The pcDNA3-Flag-RNPS1 expression plasmid was described previously^18^. To construct the expression plasmids, pcDNA3-AURKB/E5-WT and the pcDNA3-AURKB/E5-Δ#3, the fragments containing the 3′ end of intron 4, exon 5 and the 5′ end of intron 5 were amplified by

PCR from pcDNA3-AURKB/E5-E6 and pcDNA3-AURKB/E5-E6-Δ#3 plasmids^18^, respectively, and these were subcloned into pcDNA3 vector (Invitrogen–Thermo Fisher Scientific). The pcDNA3-AURKB/E5/inR-E6 and pcDNA3-AURKB/E5/inL-E6 plasmids were constructed using overlap extension PCR as previously described^31^. All PCRs were performed with PrimeSTAR Max DNA Polymerase (Takara Bio) using the described primer sets (Table S1).

### siRNA-mediated knockdown and splicing assay

The transfection of HeLa and HEK293 cells with siRNAs (100 pmol each) was performed using Lipofectamine RNAi max (Invitrogen-Thermo Fisher Scientific) according to manufacturer’s protocol. The siRNAs targeting RNPS1, ACIN1, PNN, SAP18, GPATCH8 and LUC7L3 (Table S1) were purchased (Nippon Gene).

At 72 h post-transfection, total RNAs were isolated from HeLa and HEK293 cells using the NucleoSpin RNA kit (Macherey-Nagel). To examine endogenous mRNA expression levels and the splicing products, total RNAs were reverse transcribed using PrimeScript II reverse transcriptase (Takara Bio) with oligo-dT and random primers, and the obtained cDNAs were analyzed by PCR and quantitative PCR (qPCR) using specific primer sets (Table S1).

All primers oligonucleotides were purchased (Fasmac) and all PCRs were performed with Blend Taq polymerase (Toyobo). The PCR products were analyzed by 6% polyacrylamide gel electrophoresis (PAGE). PCR products of aberrantly spliced and unspliced pre-mRNAs were quantified using NIH Image J software. Eco Real-Time PCR system (Illumina) was used for qPCR analysis with Thunderbird SYBR qPCR Mix (Toyobo). Relative changes of splicing efficiency is calculated as described previously^32^. All the experiments were independently repeated three times. Mean values, standard error of the mean (SEM), and Welch’s t-test were calculated using Excel (Microsoft).

### Western blotting analysis

At 72 h post-transfection, total protein was isolated from siRNA-treated HeLa and HEK293 cells to examine the endogenous protein expression levels. Cells were suspended in Buffer D [20 mM HEPES (pH 7.9), 50 mM KCl, 0.2 mM EDTA, 20% glycerol], sonicated for 20 sec, centrifuged to remove debris, and the lysates were subjected to Western blotting. The procedure, detection and analysis were described previously^31^.

The following antibodies were used to detect target proteins: anti-RNPS1 (1:750 dilution)^3^, anti-ACIN1 (1:1500 dilution; Cells Signaling Technology), anti-PNN (1:1500 dilution; Sigma-Aldrich), anti-SAP18 (1:1500 dilution; Proteintech), anti-AURKB (1:1500 dilution; Cells Signaling Technology) and anti-GAPDH (1:1500 dilution; MBL Life Science) antibodies.

### RNA immunoprecipitation assay

HeLa cells (in 60-mm dishes) were co-transfected with pcDNA3-Flag-RNPS1 and pcDNA3-AURKB/E5-WT or the pcDNA3-AURKB/E5-Δ#3 using Lipofectamine 2000 (Invitrogen–Thermo Fisher Scientific). After culture for 48 h, the transfected cells were suspended in 100 µL of buffer D [20 mM HEPES-KOH (pH 7.9), 50 mM KCl, 0.2 mM EDTA, 20% (v/v) glycerol] and sonicated for 20 sec (UR-20P; Tomy Seiko). The lysate was spun down to remove debris and 5 µL of supernatant was saved as 5% of input. The remaining supernatant was rocked at 4°C for 3 h with each antibody conjugated with Dynabeads Protein A (Invitrogen–Thermo Fisher Scientific) in 0.5 mL of NET2 buffer [50 mM Tris-HCl (pH7.5), 150 mM NaCl, 0.05% Nonidet P-40]. The beads were washed six times with 0.8 mL of NET2 buffer and bound RNA, together with the supernatant (5% of input), were eluted separately by TRI Reagent (Molecular Research Center). These RNAs were reverse transcribed by PrimeScript II reverse transcriptase (Takara Bio) with SP6 primer, and qPCRs were performed using specific primer sets (Table S1).

The values of % input (Figure 1D) were calculated from obtained qRT–PCR-based threshold cycle (Ct) values of immunoprecipitated RNA (Ct1) and 5% input RNA (Ct2) with the formula: 2(Ct2–Ct1) × 20.

### Whole-transcriptome sequencing (RNA-Seq) analysis

The RNA libraries were prepared from three RNA sources; HEK293 cells transfected with siRNAs targeting RNPS1 and PNN (with control siRNA). mRNA isolation, cDNA library construction and whole-transcriptome sequencing was performed by Macrogen Japan Corporation.

The obtained reads were mapped to the human genome as follows. FASTQ files were filtered by Trimmomatic v0.36^33^ with the parameters “PE-phred33 ILLUMINACLIP:Trimmomatic-0.36/adapters/TruSeq3-PE.fa:2:30:10 LEADING:3 TRAILING:3 SLIDINGWINDOW:4:15 MINLEN:150” and trimmed by fastx_trimmer included in the FASTX-Toolkit v0.0.13 (http://hannonlab.cshl.edu/fastx_toolkit/index.html) with the parameter “-l 150”. The qualified FASTQ files were mapped to human genome hg19 by TopHat v2.1.1^34^ and the transcripts were assembled by Cufflinks v2.2.1^35^ with default parameters. PSI (percent-spliced-in) values were estimated by MISO v0.5.2^36^ using the assembled transcripts as previously described^37^. RNA-Seq raw data (see Figures 2A, S4) have been deposited in DDBJ database (http://www.ddbj.nig.ac.jp/index-e.html) under accession No. DRA16802.

### Cell cycle synchronization experiments

To achieve cell synchronization at G2/M phase, HEK293 cells were plated (in 35-mm dishes) at 40% confluency. Cells were subsequently treated with 2 mM thymidine (Sigma-Aldrich) for 24 h, washed with PBS, and supplemented with fresh DMEM. After culture for 3 h, cells were followed by 12 h culture in the presence of microtubule-disrupting agent nocodazole at 100 µM concentration (Sigma-Aldrich). To remove nocodazole, cells were washed with phosphate-buffered saline (PBS) and cultured in fresh DMEM. Cells were harvested every 4 h for 12 h after the removal of nocodazole.

To achieve cell synchronization at G1/S phase, HEK293 cells were plated (in 35 mm dishes) at 20% confluency. Cells were treated with 2 mM thymidine for 18 h, washed with PBS, and supplemented with DMEM. After culture for 9 h, cells were treated with 2 mM thymidine for another 15 h. Cells were released from thymidine block by washing with PBS followed by culture in fresh DMEM. Cells were harvested every 4 h for 12 h after the removal of thymidine. To inhibit proteasome, HEK293 cells were treated with MG132 (Sigma-Aldrich) at a final concentration of 1 µM. After 4 h and 8 h of culture, cells were harvested and total protein was isolated (see above).

### Quantification and statistical analysis

Three independent experiments were performed. Mean values, SE values, and Welch’s t-test were calculated using Excel (Microsoft).

## ACKNOWLEDGEMENTS

We thank H. Shirasaki and T. Kanehisa for technical support, and members of our labs for constructive discussions. K. Fukumura was partly supported by Grants-in-Aid for Scientific Research (C) and Challenging Research (Exploratory) [Grant numbers: JP18K07304, JP21K07202, JP24K21953] from the Japan Society for the Promotion of Science (JSPS), a Research Grant from the Hori Sciences and Arts Foundation, a Research Grant from Nitto Foundation, a Research Grant from Takeda Science foundation, a Research Grant from Aichi Cancer Research Foundation, and a Research Grant from the Mochida Memorial Foundation. A. Masuda was partly supported by Grants-in-Aid for Scientific Research (B) and Challenging Research (Exploratory) [Grant numbers: JP21H02476, JP21H02838, JP22K19269] from JSPS, and a Grant from the Japan Agency for Medical Research and Development [Grant number: JP23ek0109497]. K. Ohno was partly supported by Grants-in-Aid for Scientific Research (B) [Grant number: JP23H02794] from JSPS and Health and Labour Science Research Grant [Grant number: JP23FC1014] from the Ministry of Health Labour and Welfare. A. Mayeda was partly supported by Grants-in-Aid for Scientific Research (B) and Scientific Research (C) [Grant number: JP16H04705, JP21K06024] from JSPS. A. Mayeda was awarded a Professor Emeritus of the Fujita Health University in April 2024.

## AUTHOR CONTRIBUTIONS

K. Fukumura and A. Mayeda conceived and designed the experiments; K. Fukumura performed most of the experiments and analyses, organized the data and drafted the manuscript; J.-i. Takeda and A. Masuda performed bioinformatics analyses of the sequencing data; K. Fukumura, K. Ohno and A. Mayeda revised and edited the manuscript; and O. Nagano and H. Saya for sharing facility and financial support. A. Mayeda coordinated and supervised the whole project. All authors read, corrected and approved the final manuscript.

---

**Table S1.**
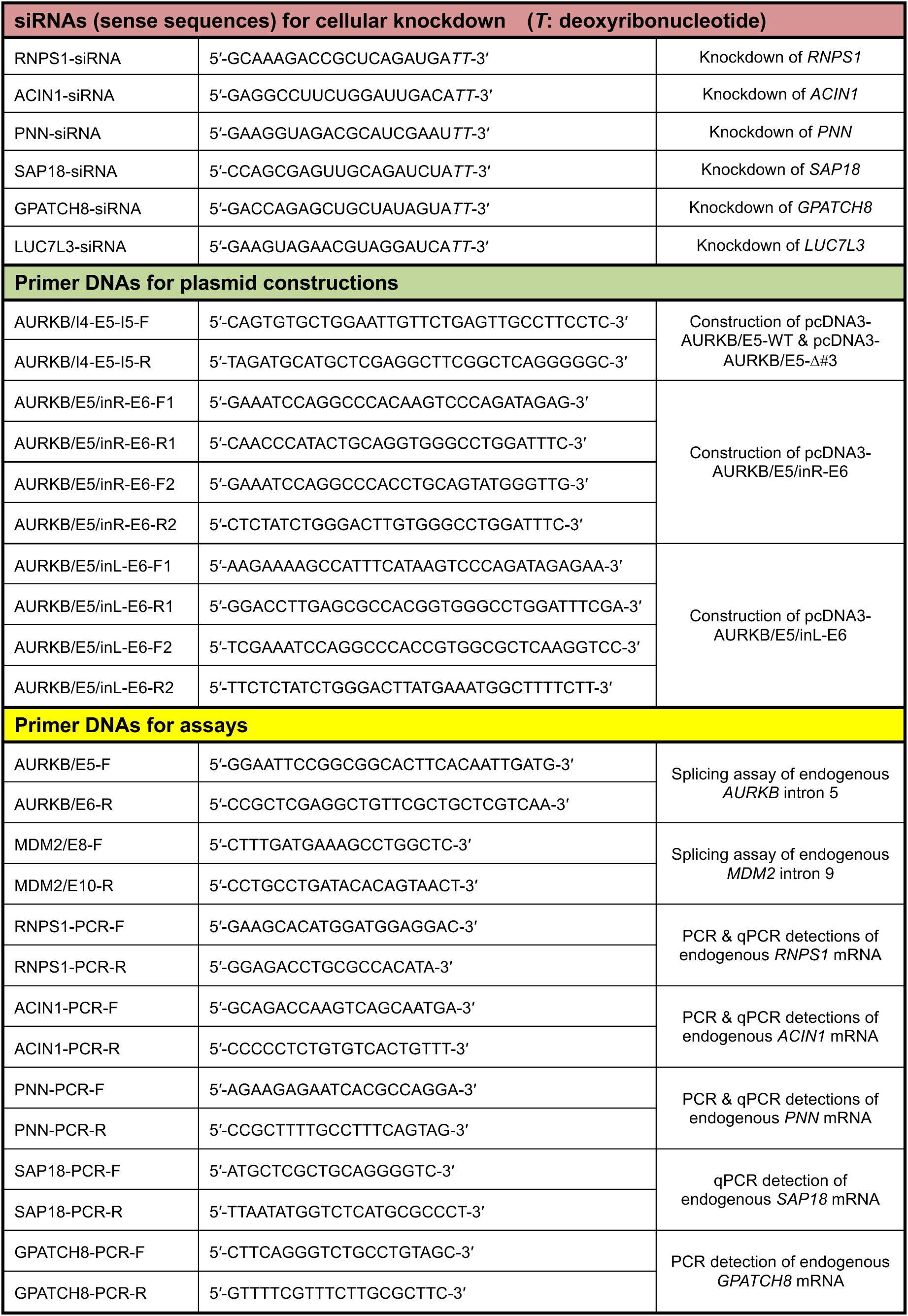
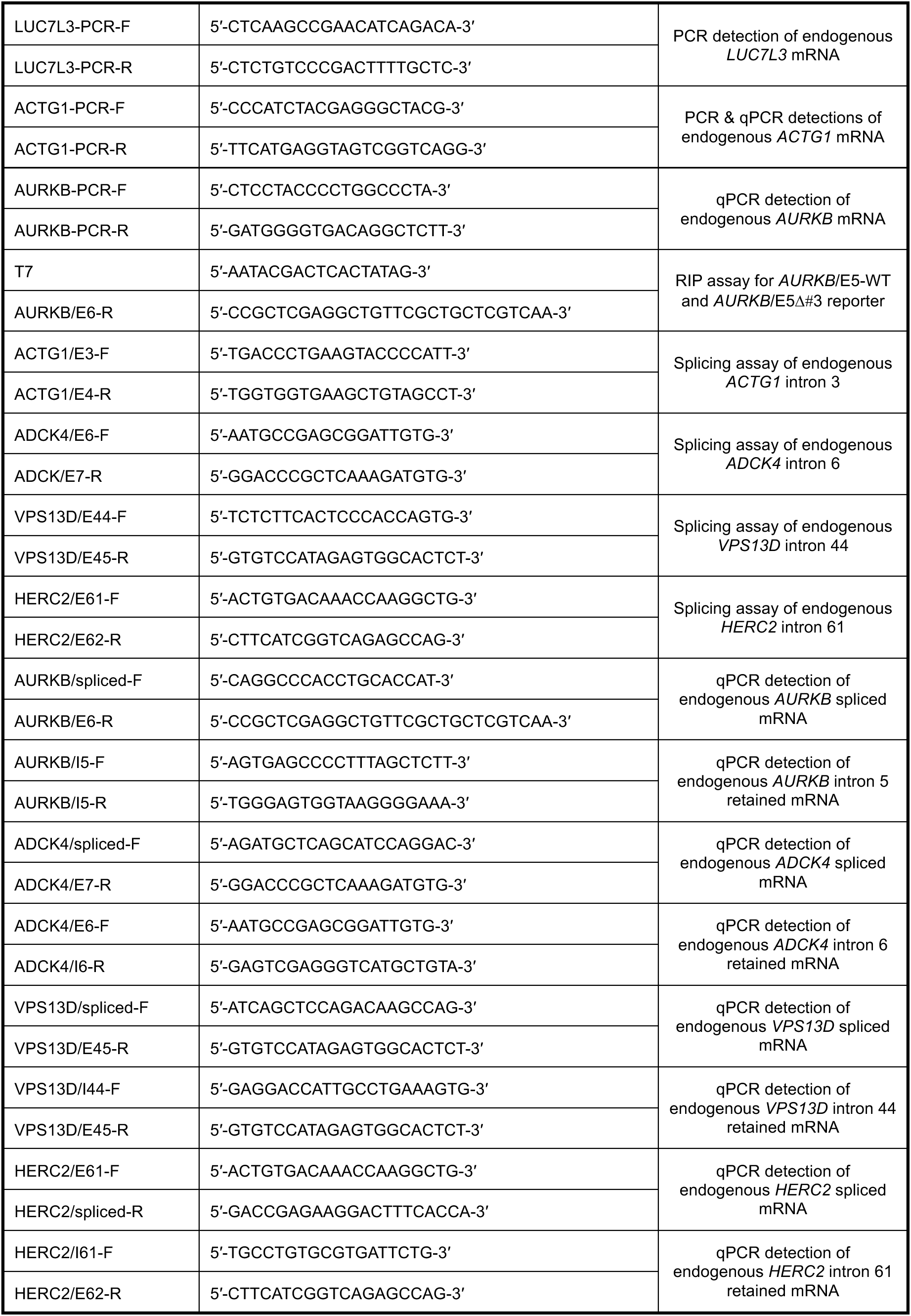
List of all the synthetic oligonucleotides used in the experiments.

**Figure S1.**
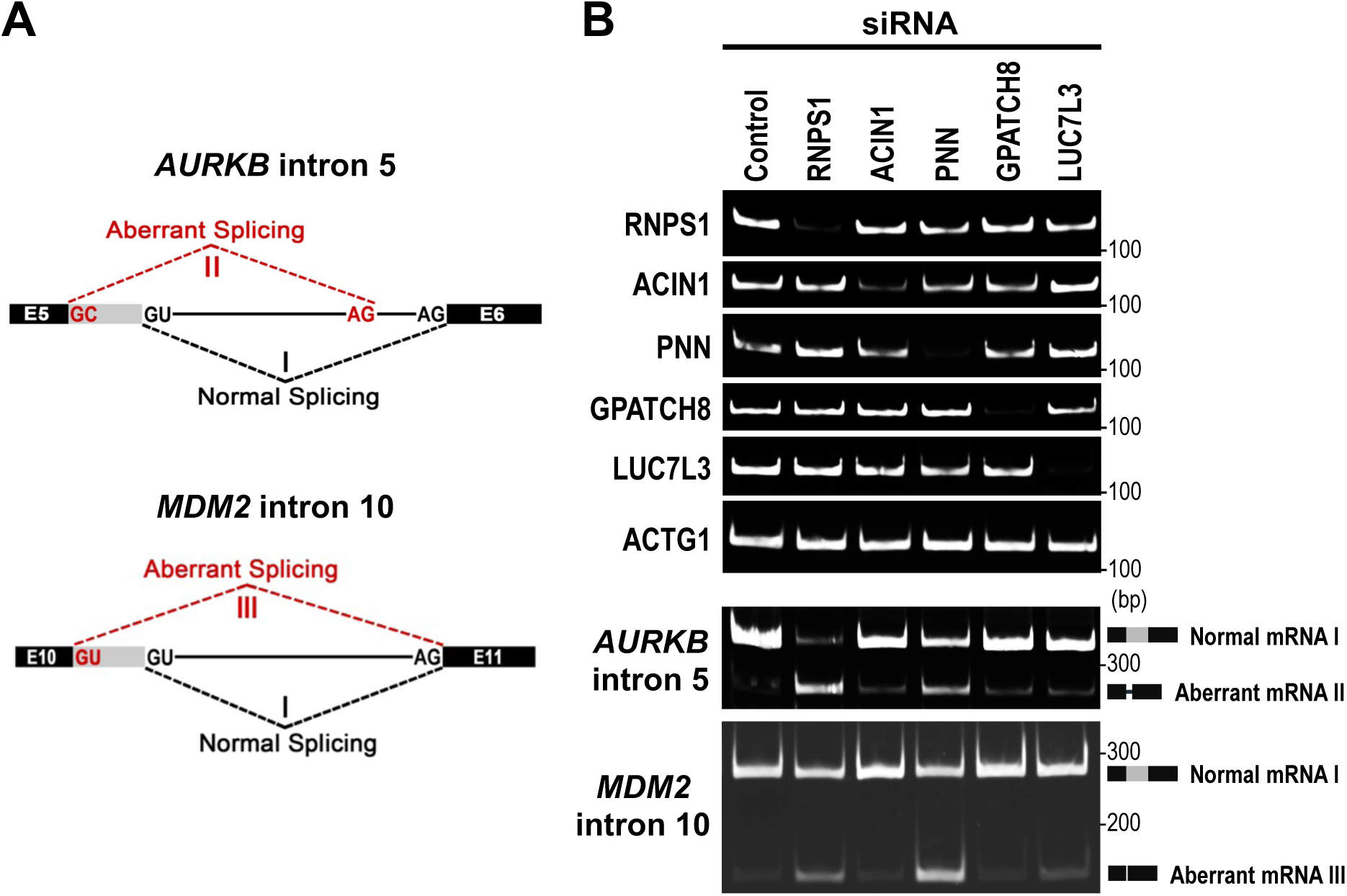
PSAP components, but not ASAP-specific component and other RNPS1-binding factors, promote normal splicing of two PSAP-controlled pre-mRNAs (Related to Figure 1C) **(A)** Schematic diagram of pre-mRNA splicing pathways that generate the normal mRNA (I) and aberrant mRNAs (II, III) in the indicated pre-mRNA. Previously it was shown that these aberrant mRNAs (II, III) are generated by siRNA-mediated depletion of RNPS1, a component of of PSAP [S1]. **(B)** siRNA-mediated depletion of the ASAP-specific component (ACIN1), PSAP-specific component (PNN), and other RNPS1-binding proteins (GTATCH8 and LUC7L3) in HeLa cells and these proteins were analyzed by Western blotting with indicated antibodies (upper panel). Indicated spliced mRNAs (I, II, III) from the endogenous *AURKB* and *MDM2* genes were detected by RT–PCR and visualized by PAGE (lower panel). *GPATCH8* (G patch domain-containing protein 8) was isolated from human brain cDNA library and its function is unknown. Previous exome sequencing study showed that a mutation in *GPATCH8* associated with hyperuricemia co-segregating with osteogenesis imperfecta [R2]. *LUC7L3* (LUC7 like3) is a paralogues of yeast U1 snRNP component *LUC7*, which is involved in alternative splicing regulation in human [S3, S4].

**Figure S2.**
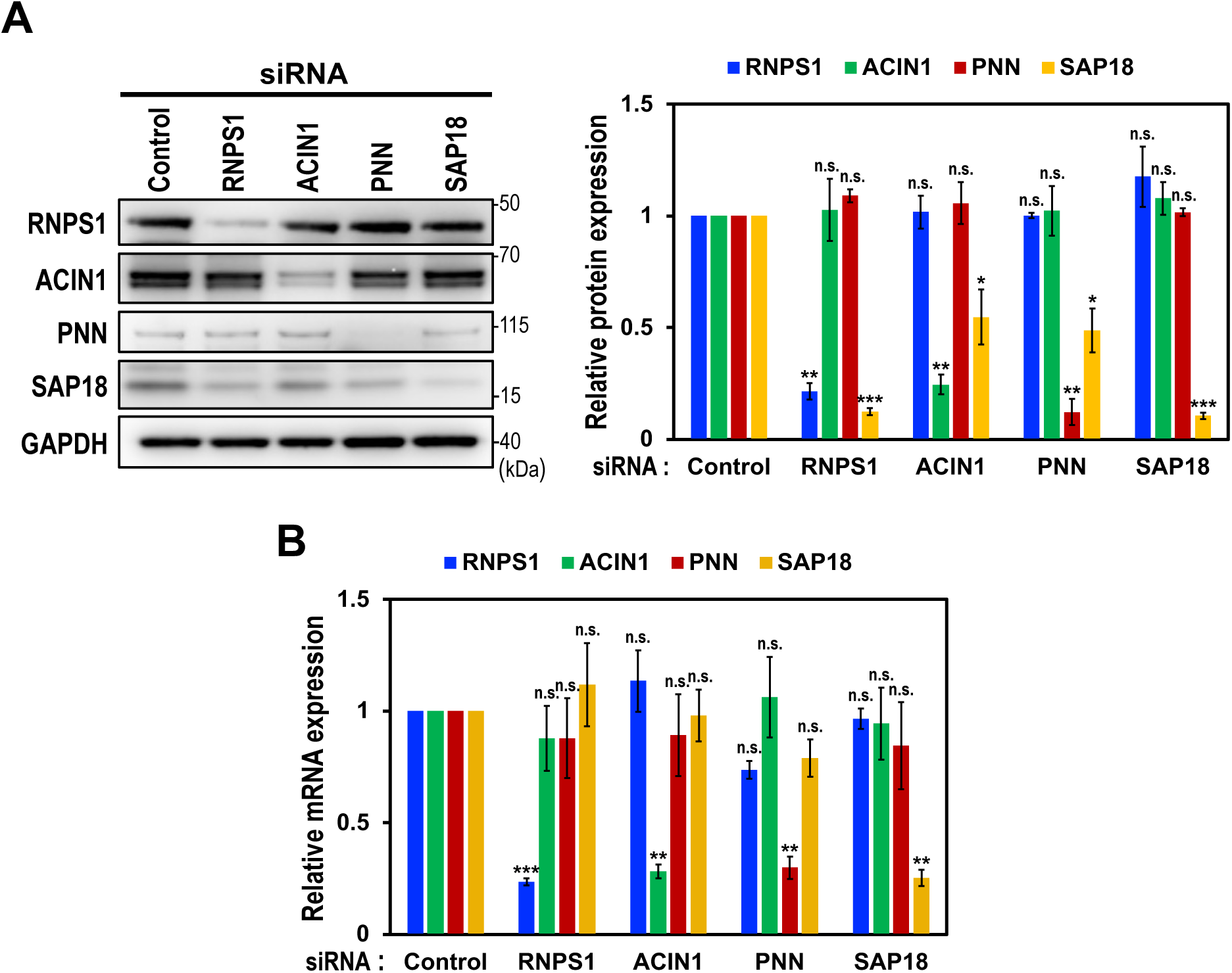
Knockdown of RNPS1, ACIN1 and PNN in HEK293 cells also caused co-depletion of SAP18 protein (Related to Figure 1A, B) **(A)** siRNA-mediated depletion of the indicated ASAP/PSAP components in HEK293 cells were analyzed by Western blotting with indicated antibodies (left panel). Individual bands on the Western blots were quantified and the relative values were standardized to that in the control siRNA (right graph). Means ± standard errors (SE) are given for three independent experiments and Welch’s t-test values were calculated (*p < 0.05, **p < 0.005, ***p < 0.0005, n.s. p > 0.05). **(B)** The mRNA levels of ASAP/PSAP components in (A) were analyzed by RT-qPCR using specific primer sets. See (A) for the statistical analysis.

**Figure S3.**
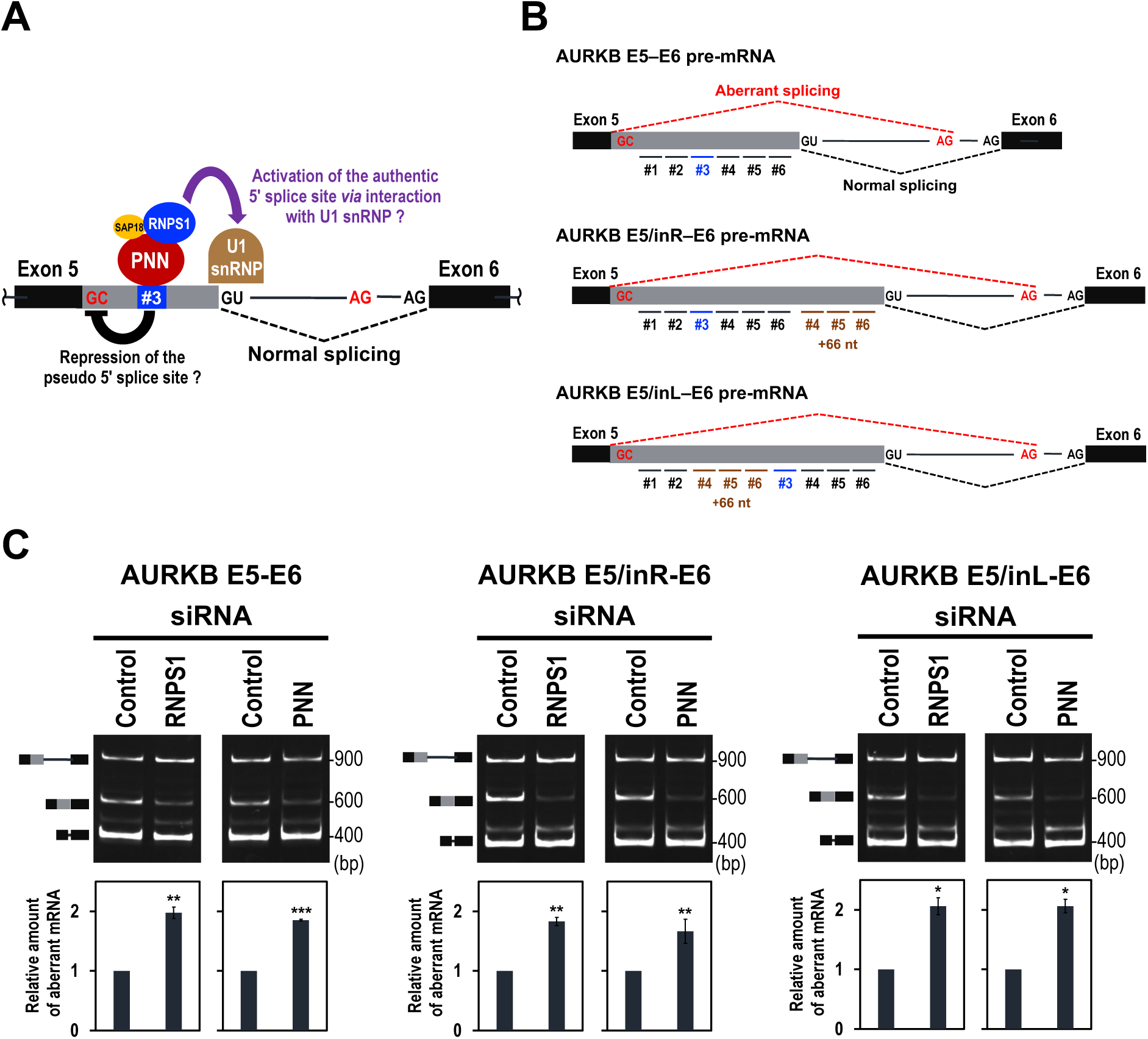
PSAP either represses the upstream pseudo 5′ splice site or activates the downstream authentic 5′ splice site to ensure precise splicing of *AURKB* pre-mRNA (Related to Figure 1C) **(A)** Schematic diagram of two conceivable PSAP-mediated mechanisms to maintain normal splicing. (i) PSAP bound on the target site (#3) represses the upstream pseudo 5’ splice site through either direct masking or indirect interference. (ii) PSAP on the target site (#3) activates the downstream authentic 5’ splice site through RNPS1 interaction with a component of U1 snRNP [S1]. **(B)** Three *AURKB* mini-gene transcripts to test the above models. The distances from PSAP-binding site (#3) to either upstream authentic 5’ splice site or downstream pseudo 5’ splice site were expanded by inserted duplicated fragments (#4-#5-#6) in E5/inR–E6 and E5/inL–E6 mini-genes, respectively. **(C)** *In cellulo* splicing assay of ectopically expressed these three *AURKB* mini-genes upon depletion of PSAP components, RNPS1 and PNN. Splicing products were detected by RT– PCR, visualized by PAGE, and individual bands on the PAGE gel were quantified. See the legend to Figure 1C for the methods of quantification and statistical analysis.

**Figure S4.**
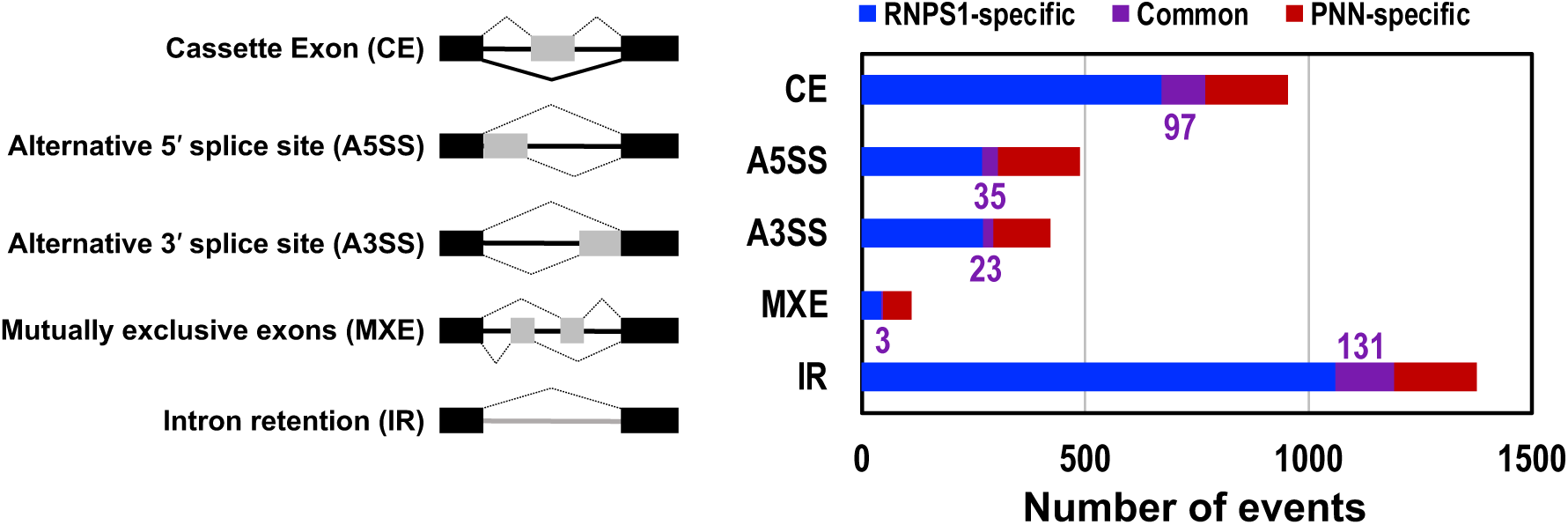
Knockdown of PSAP components, RNPS1 and PNN, causes global changes in splicing patterns (Related to Figure 2A) RNA-Seq analyses of RNPS1- and PNN-knockdown HEK293 cells were performed. The number of statistically significant changes in each alternative splicing pattern (left panel) is visualized as bar graph (right graph). The purple numbers show the common splicing changes, in either directions, induced in RNPS1- and PNN-knockdown HEK293 cells.

**Figure S5.**
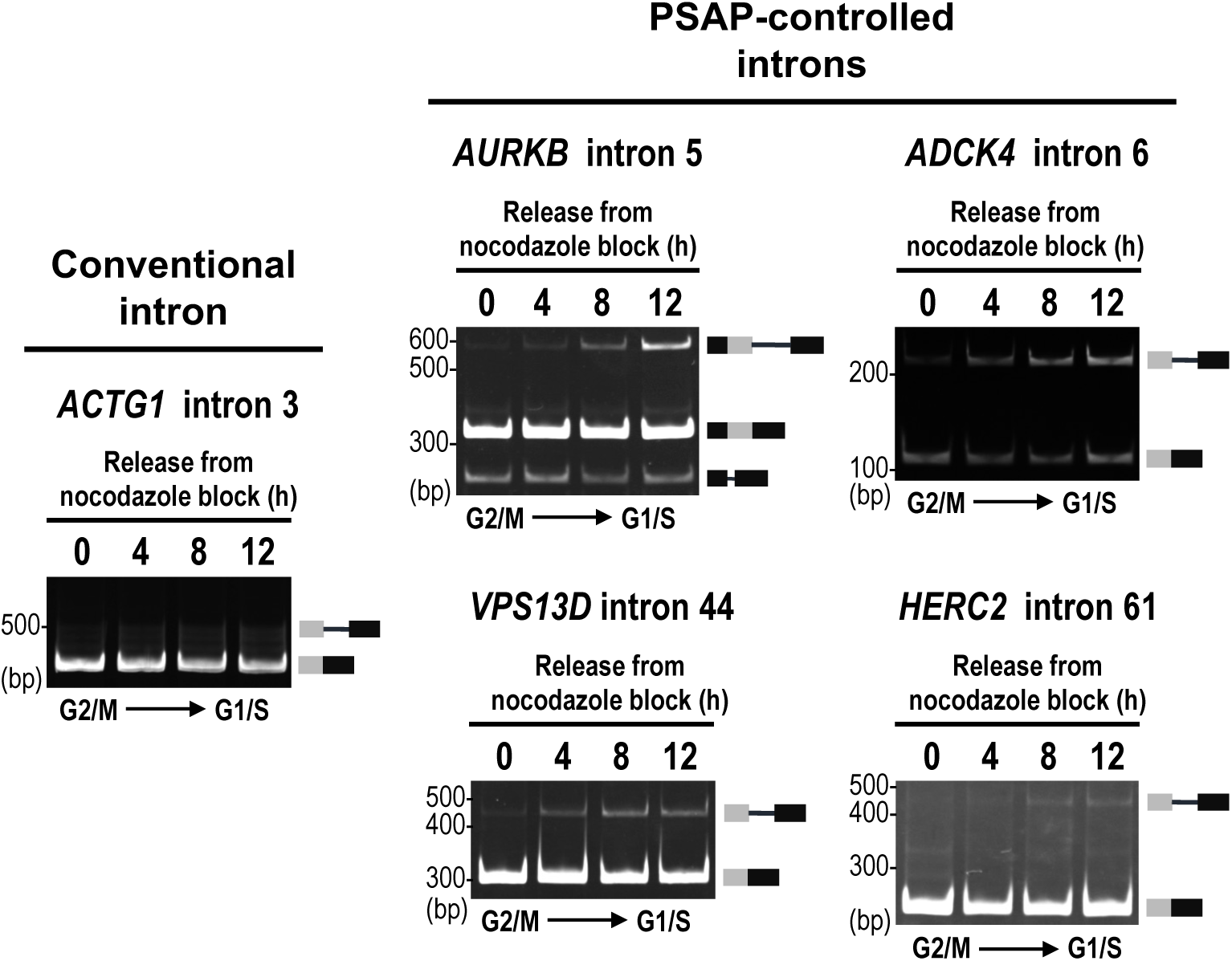
Periodic generation of unspliced introns from G2/M to G1/S phase were observed in PSAP-controlled introns but not in conventional intron (Related to Figures 3B, 4) HEK293 cells were synchronized in G2/M phase by nocodazole block. Endogenous splicing in the indicated conventional intron and PSAP-controlled introns was analyzed by RT-PCR followed by PAGE.

**Figure S6.**
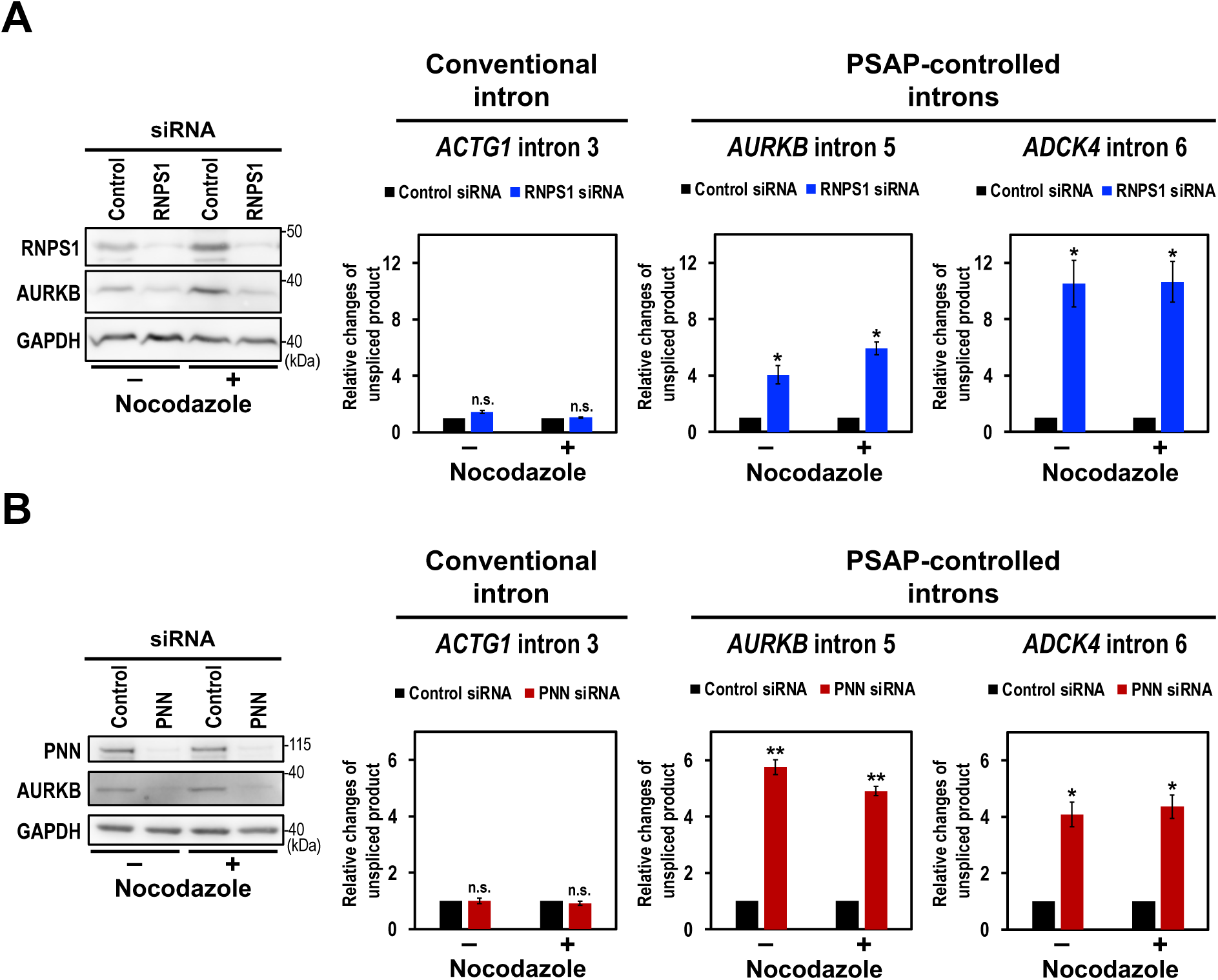
Knockdown of RNPS1 and PNN generates unspliced products in PSAP-regulated introns, but not in conventional intron, irrespective of cell cycle phase (Related to Figures 3–5) **(A)** RNPS1 was depleted by siRNA-mediated knockdown in HEK293 cells and treated with nocodazole to arrest the cell cycle at the G2/M phase. Proteins were analyzed by Western blotting with indicated antibodies. *AURKB* is downregulated by RNPS1-knockdown as previously described [S1]. Indicated retained introns were analyzed by RT–qPCR and the values were standardized to that of the control-knockdown and plotted. Means ± SE are given for three independent experiments and Welch’s t test values were calculated (*p < 0.05, n.s. p > 0.05). **(B)** PNN was depleted by siRNA-mediated knockdown in HEK293 cells and treated with nocodazole to arrest the cell cycle at the G2/M phase. The same analyses as in (A) were performed. See (A) for the statistical analysis (*p < 0.05, **p < 0.005, n.s. p > 0.05).

